# Dynamic representation of multidimensional object properties in the human brain

**DOI:** 10.1101/2023.09.08.556679

**Authors:** Lina Teichmann, Martin N. Hebart, Chris I. Baker

**Affiliations:** Laboratory of Brain and Cognition, National Institute of Mental Health, National Institutes of Health, Bethesda MD, USA; Vision and Computational Cognition Group, Max Planck Institute for Human Cognitive and Brain Sciences, Leipzig, Germany; Department of Medicine, Justus Liebig University Giessen, Giessen, Germany; Center for Mind, Brain and Behavior (CMBB), Universities of Marburg, Giessen, and Darmstadt, Germany

## Abstract

Our visual world consists of an immense number of unique objects and yet, we are easily able to identify, distinguish, interact, and reason about the things we see within a few hundred milliseconds. This requires that we integrate and focus on a wide array of object properties to support diverse behavioral goals. In the current study, we used a large-scale and comprehensively sampled stimulus set and developed an analysis approach to determine if we could capture how rich, multidimensional object representations unfold over time in the human brain. We modelled time-resolved MEG signals evoked by viewing single presentations of tens of thousands of object images based on millions of behavioral judgments. Extracting behavior-derived object dimensions from similarity judgments, we developed a data-driven approach to guide our understanding of the neural representation of the object space and found that every dimension is reflected in the neural signal. Studying the temporal profiles for different object dimensions we found that the time courses fell into two broad types, with either a distinct and early peak (∼125 ms) or a slow rise to a late peak (∼300 ms). Further, early effects were stable across participants, in contrast to later effects which showed more variability, suggesting that early peaks may carry stimulus-specific and later peaks more participant-specific information. Dimensions with early peaks appeared to be primarily visual dimensions and those with later peaks more conceptual, suggesting that conceptual representations are more variable across people. Together, these data provide a comprehensive account of how behavior-derived object properties unfold in the human brain and form the basis for the rich nature of object vision.

## Introduction

A central aspect of vision is our ability to understand and interact with a huge variety of different objects that are associated with a wide range of perceptual (e.g., color, shape), functional (e.g., affordances), and conceptual (e.g., associations) properties. The importance of each of these properties may even vary depending on context and behavioral goals. What neural representations support our ability to make sense of our visual world in the span of just a few hundred milliseconds? To capture the multidimensional nature of object representations and how they unfold over time, here we moved beyond the traditional, stimulus-focused approach of studying object processing by directly examining how continuous, behavior-derived object dimensions are reflected in the neural signal.

Prior studies that have typically focused on specific types of stimuli or properties, have demonstrated both univariate (e.g., N170 for faces) and multivariate differences in the brain response over time. While these studies have revealed some general features of the object response, they are often based on relatively small, hand-selected sets of stimuli (e.g., Bankson et al., 2018; Carlson et al., 2013a; Cichy et al., 2014) which do not sample the object space in a representative way and cannot adequately capture the richness of the object response (Grootswagers & Robinson, 2021). Hand-selecting sets of stimuli may lead to a sampling bias because some types of objects are considered to be important and may be overrepresented (e.g., faces, animals) while others may be completely absent or underrepresented (e.g., furniture, cars, birds).

To provide a more comprehensive understanding of object vision, we focused on addressing two key challenges: 1) sampling an expansive set of object stimuli across the thousands of object types we can identify and interact with, and 2) accounting for the rich meaning and behavioral relevance associated with individual objects beyond discrete labels (Contier et al., 2024). To do this, we turned to THINGS-data (Hebart, Contier, Teichmann et al., 2023), which contains MEG data for 1,854 systematically-sampled object concepts (Hebart et al., 2019) as well as rich behavioral data comprising 4.7 million similarity judgments that have been used to derive 66 underlying dimensions of objects in a data-driven manner (e.g., colorful, plant-related, transportation-related) (Hebart et al., 2020, Hebart, Contier, Teichmann et al., 2023). Using these data, we developed a data-driven approach to uncover the temporal dynamics of object processing by directly examining how behavior-derived object dimensions are reflected in the evolving object representations in the human brain.

In contrast to previous work requiring category-based stimulus selection and labelling, here we use behavioral embeddings that characterize each image across multiple dimensions capturing similarity relationships directly. In addition, the dimension values for each object are continuous (e.g., jellybeans are more colorful than sunflowers, but sunflowers are more colorful than sugar), allowing for fine-grained modelling of similarity in the neural data and thus capturing the richness of object vision. In contrast to common approaches such as representational similarity analysis (RSA), our method allows us to directly study evoked neural representations at the global level (i.e., across all dimensions) as well as at the local level (i.e., each dimension separately). Critically, our approach goes beyond studying object identification and categorization and allows us to determine the time course of response specific to each of the behavior-derived dimensions.

Our results show that every dimension is reflected in the neural signal. The temporal profiles evoked by the dimensions tended to group according to the relative strength of two phases of processing (∼125 ms and ∼300 ms) as well as the presence or absence of an offset related response (∼500-600 ms). Critically, early effects were more generalizable across participants while later effects were more variable across people. This suggests that stimulus-specific information is reflected in the early parts of the signal while subject-specific information unfolds later in time. An exception to this were dimensions that are primarily associated with physical properties of the object, which generalized well across participants throughout the timeseries. By focusing on behavioral relevance of object properties, our results collectively provide a comprehensive characterization of the temporal unfolding of visual object responses in the human brain.

## Results

The overarching goal of the current study was to characterize how multidimensional representations unfold over time by combining large-scale MEG data with behavior-derived similarity embeddings. Our primary aims were to (1) extract time courses from the MEG response that are associated with each behavior-derived multidimensional profile, (2) reveal how these time courses vary across dimensions and participants, and (3) identify prototypical temporal characteristics shared between response profiles of individual dimensions.

THINGS-MEG measured evoked neural responses in four participants viewing >27,000 unique natural images associated with 1,854 object concepts. To associate object dimensions with these natural images, we used behavioral embeddings derived from similarity judgments, based on 4.7 million judgments on 1,854 object concepts in a triplet odd-one out task (Hebart, Contier, Teichmann et al., 2023). Thus, the stimuli used in the MEG study are associated with both object concept labels (e.g., nail polish) as well as weights on behavior-derived dimensions (e.g., colorful) (Hebart et al., 2020). The dimensions cover a broad range of object properties, with some being strongly linked to visual features (e.g., colorfulness) and others linked more to functional or contextual features (e.g., childhood-related). The behavioral similarity embeddings were based on concept-level judgments (i.e., one image per object concept), potentially missing some of the visual variability in the MEG stimuli. To overcome this issue, we used an artificial neural network (CLIP-VIT, Radford et al., 2021) to augment the behavioral dataset and generate image-level embeddings for later predictions (Hebart et al., 2022). Post-hoc analyses showed a consistent improvement in prediction scores for all 66 dimensions when using image-level versus concept-level embeddings. While the results were consistently stronger, the overall pattern of results remained similar even without the use of CLIP-VIT, specifically for more semantic dimensions.

We used the scores on the 66 behavior-derived dimensions to model the evoked neural response to >26,000 images recorded with MEG. In particular, we used both decoding and encoding models to associate the multivariate MEG-sensor response with the 66-dimensional behavioral embedding. In contrast to previous work, our method allows us to effectively examine *multidimensional* object profiles that are obtained from behavioral data in a data-driven way. We can capture and examine the relationships between objects across many dimensions, as two images that are similar along one dimension may be very different along another dimension (e.g., beads and nail polish are both colorful but not necessarily childhood-related). Thus, this approach does not rely on selecting and contrasting object classes but instead uses the same images and experimental trials with a relabeling according to behavior-derived dimensional profiles.

### (1) Distinct time courses can be derived for each behavior-derived dimension

We fitted multiple linear regression models to learn the association between the MEG sensor activation patterns and the behavioral embeddings at every timepoint (Figure 1). The linear models were fit on MEG data from 11 sessions (20,394 trials). Using the left-out, independent session (1,854 trials) as a test set, we predicted the continuous value along each dimension from the MEG sensor activation patterns. Correlating these predicted scores with the true behavioral dimensional profiles resulted in timeseries of dimension information in the neural response for all four participants. The analysis revealed behavior-derived multidimensional information from ∼80 ms onwards (Figure 2A). The time course showed an early peak at ∼100 ms which was maintained over time up to 1000 ms after stimulus onset, with an overall peak at ∼ 300ms. To examine the relevance of individual MEG sensors to this effect, we also fitted linear models to predict each sensor activity pattern separately. In particular, we trained models to predict the univariate response in each MEG sensor at every timepoint using the multidimensional behavioral values associated with each stimulus. Our results showed that, while posterior sensors had the strongest effect across time, the relative contribution of frontal sensors increased later in time (>150 ms).

**Figure 1.**
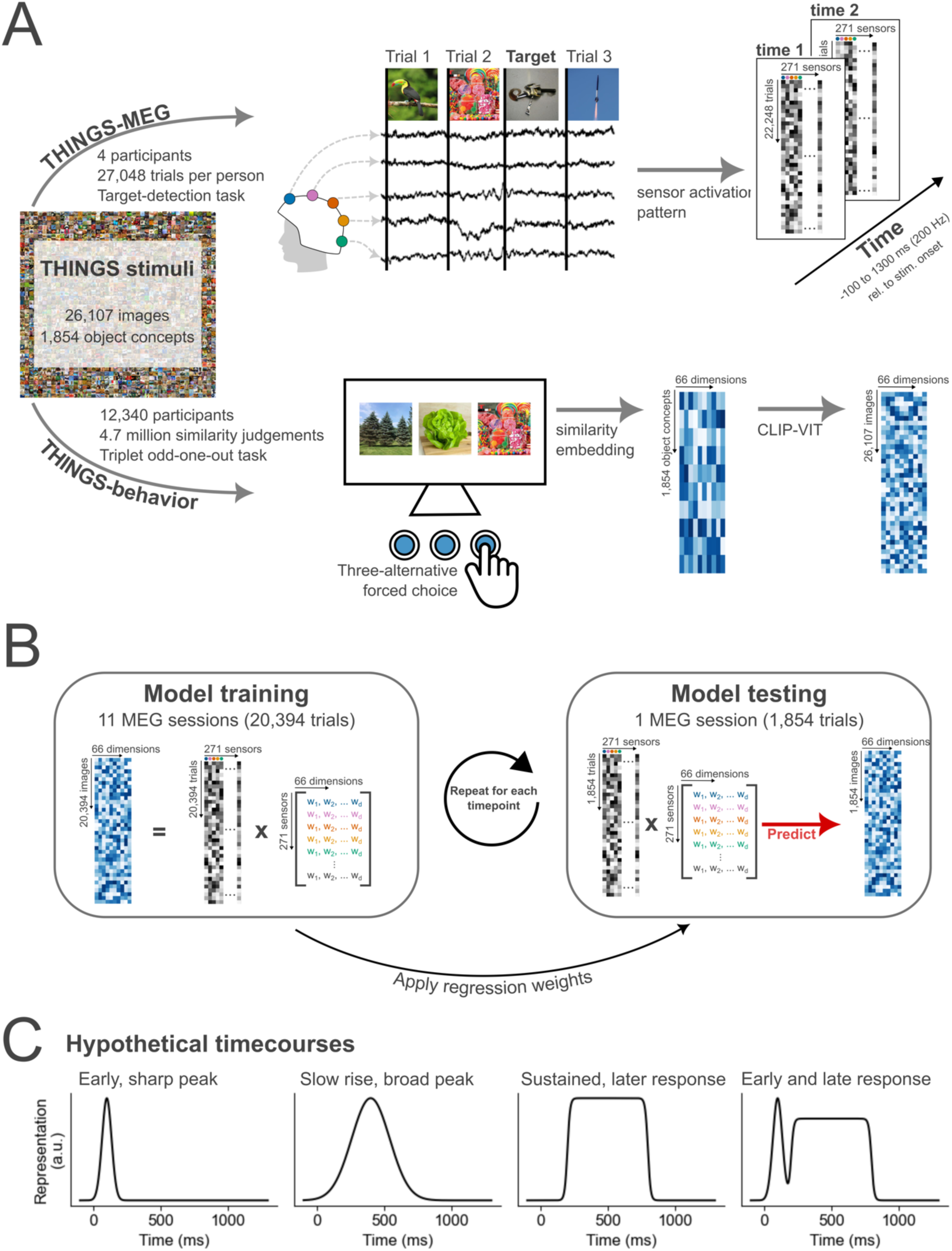
Summary of THINGS-MEG and THINGS-Behavior datasets and the methodological approach to combine them. (A) Summary of the datasets used. Evoked responses to images from the THINGS image-database were recorded over time using MEG. In total, four participants completed 12 sessions, resulting in >100,000 trials in total. During the MEG session, participants were asked to detect computer-generated images of non-nameable objects. In the behavioral task, a separate set of participants viewed three objects from the THINGS image-database at the time and were asked to pick the odd-one-out. A computational model was then trained to extract similarities along 66 dimensions for all object concepts. Using CLIP-VIT, we extended the embedding to capture similarities for every image. The data for behavioral data was crowdsourced via Amazon Mechanical Turk. In total >12,000 participants completed a total of 4.7 million similarity judgements. (B) Overview of the methodological approach of combining these two datasets with the goal of understanding how multidimensional object properties unfold in the human brain. To train the model, we extract the sensor activation pattern at each timepoint across the MEG sensors and use the behavioral embeddings to learn an association between the two datasets. The linear regression weights are then applied to sensor activation patterns of independent data to predict the behavioral embedding. To evaluate the model’s performance, we correlated the predicted and true embedding scores. (C) Hypothetical time courses that could be observed for different dimensions.

**Figure 2.**
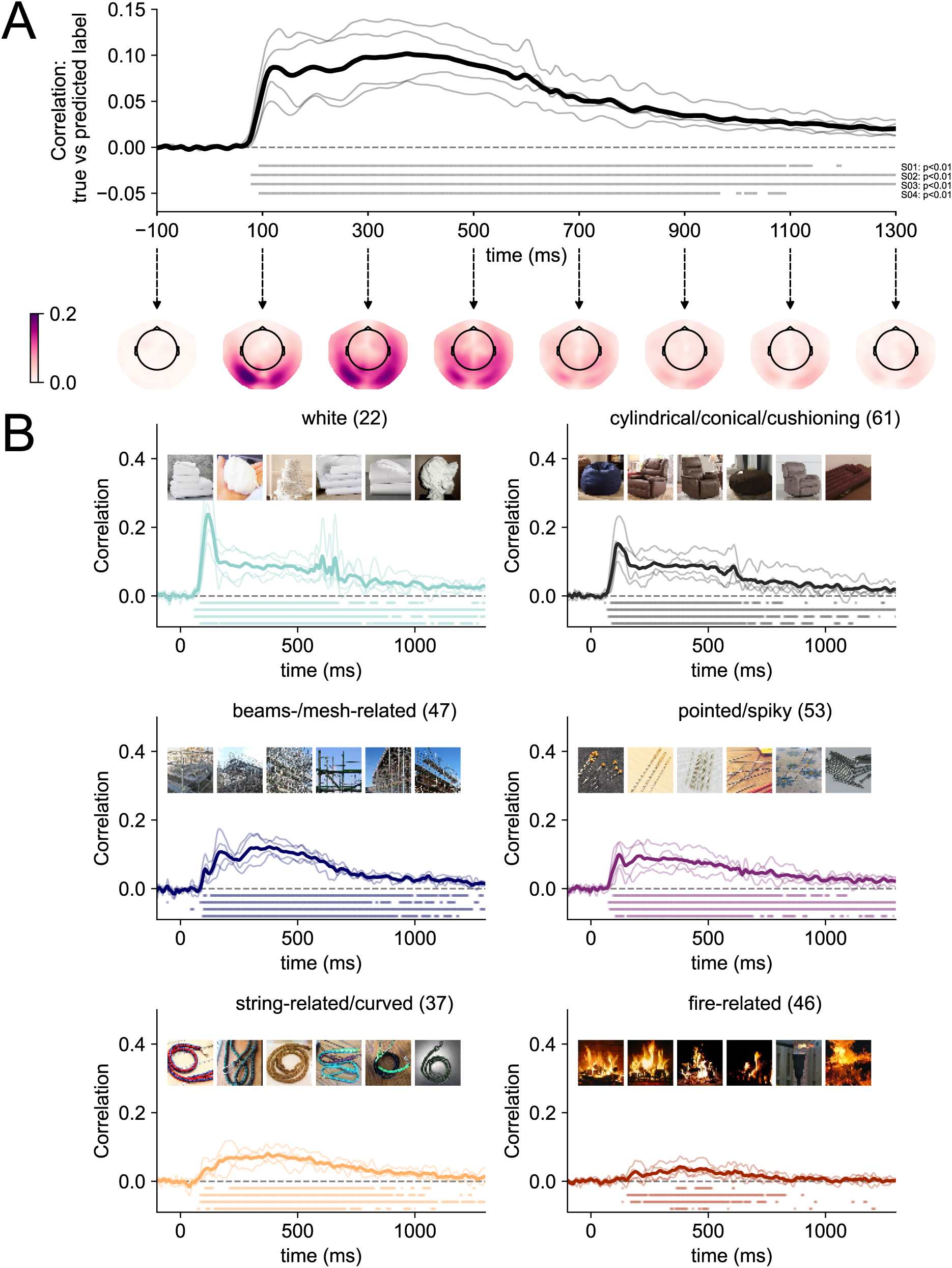
Modelling results for within-participant models of MEG data and multidimensional similarity judgments. (A) Correlation between the predicted and true behavioral embeddings across all dimensions over time. The thick, black line shows the average across all participants, the thin grey lines individual participants. The dots below the time courses show significant timepoints for each of the four participants at a threshold of p < 0.01 established based on individual-level null distributions generated using label permutations (see Methods). The topographical maps below the time courses show single timepoints of the model performance when it is fitted across all dimensions to individual sensors. Darker colors show a higher correlation between the predicted and true weights at each sensor location. A dynamic version of the topographical plots can be accessed <u>here</u>. (B) Example time courses for six dimensions. Time courses were first sorted by peak amplitude, and then we picked every eleventh time course to show a representative sample of time courses with different signal-to-noise ratios. The images within each subplot show the six stimuli with the highest weight on that dimension (derived from behavior). All individual time courses can be found in the Supplementary Materials (Figure S1). A dynamic version of the multidimensional unfolding across all 66 dimensions can be accessed <u>here</u>.

Extracting the correlation time courses for each dimension separately, we found that there was significant information about most dimensions in the signal from 100 ms onwards (for at least 3 participants, 56/66 dimensions carried significant information by 100 ms and 65/66 dimensions by 150 ms). Furthermore, our results revealed a different unfolding of neural responses across time for different dimensions. For example, dimensions such as “plant-related”, “colorful/playful”, “white” showed distinct, early peaks (∼125 ms). In contrast, other dimensions such as “body-/people-related”, “food-related”, and “transportation-related” yielded a slower rise to a later peak (∼300 ms). In addition, several of the dimensions yielding distinct early peaks exhibited a stimulus offset effect at ∼500 ms. In contrast, several other dimensions did not show a distinct early peak or offset response but unfolded slowly over time and rose to a late peak (>300 ms). While signal-to-noise ratio differed across dimensions, all dimension time courses exceeded zero at some point over time (see Figure 2B for representative example time courses selected based on peak amplitude and Supplementary Figure 1 for all time courses). Strikingly, the behavior-derived dimensional profiles were evident in the neural response even though MEG participants completed an orthogonal detection task. This demonstrates that the dimensions are automatically reflected in the neural data without a task that requires participants to engage with the object properties directly. Overall, these results highlight that a wide range of behavior-derived dimensions are reflected in distinct temporal profiles and that their information is distributed across MEG sensors.

### (2) Time courses vary across dimensions and are consistent across participants

Building on the finding that a range of behavior-derived dimensions are reflected in distinct neural profiles measured with MEG, we next tested to what degree these time courses are consistent across participants. We used multiple linear regression models (see (1)), but this time with a subject-based cross-validation scheme, to examine whether the temporal characteristics we found for each dimension are idiosyncratic or consistent across participants. Specifically, we trained the model on MEG data from each participant and tested its performance on the data from the remaining ones. We used a session-wise cross-validation scheme, so that the model was trained and tested on the same number of trials as the within-participant analysis (see section (1)). This is a very stringent test for generalizability, as the model trains and tests on completely different datasets. Our results show that for most dimensions the across-participant models revealed similar timeseries characteristics as the within-participant model (Figure 3), highlighting that the temporal profiles we uncovered for each dimension were robust and not idiosyncratic to specific individuals of our study. As Figure 3A shows, the early peaks (∼125 ms) in particular were consistent in amplitude and timing when comparing the within- and the across-participant model. In contrast, later effects (e.g., 200 ms, 400 ms, 600 ms) did not generalize as well across participants for most dimensions. For many dimensions, we observed a substantial drop in performance for the across-participant model performance relative to the within-participant model at around 200 ms before the performance improved again (Figure 3B & 3C). This indicates that the differences we observed between the within- and across model were not solely driven by a time-dependent decrease in signal-to-noise ratio. The magnitude of the differences between the within- and across-participant models were stable after the initial drop from around 250 ms onwards (Figure 3C). In addition, the results show that strong stimulus-offset effect at 500-600 ms observed in some dimensions for the within-participant models (e.g., the color dimensions) were also present when the model was trained and tested across participants. Together, these findings suggest that early effects (∼125 ms) may carry largely stimulus-specific information that generalizes well across participants, while slightly later effects (∼200 ms) are more subject-specific. However, for dimensions that are visually more homogenous (e.g., white, colorful), we found that the within- and across-models perform similarly throughout time, including the response associated with stimulus offset.

**Figure 3.**
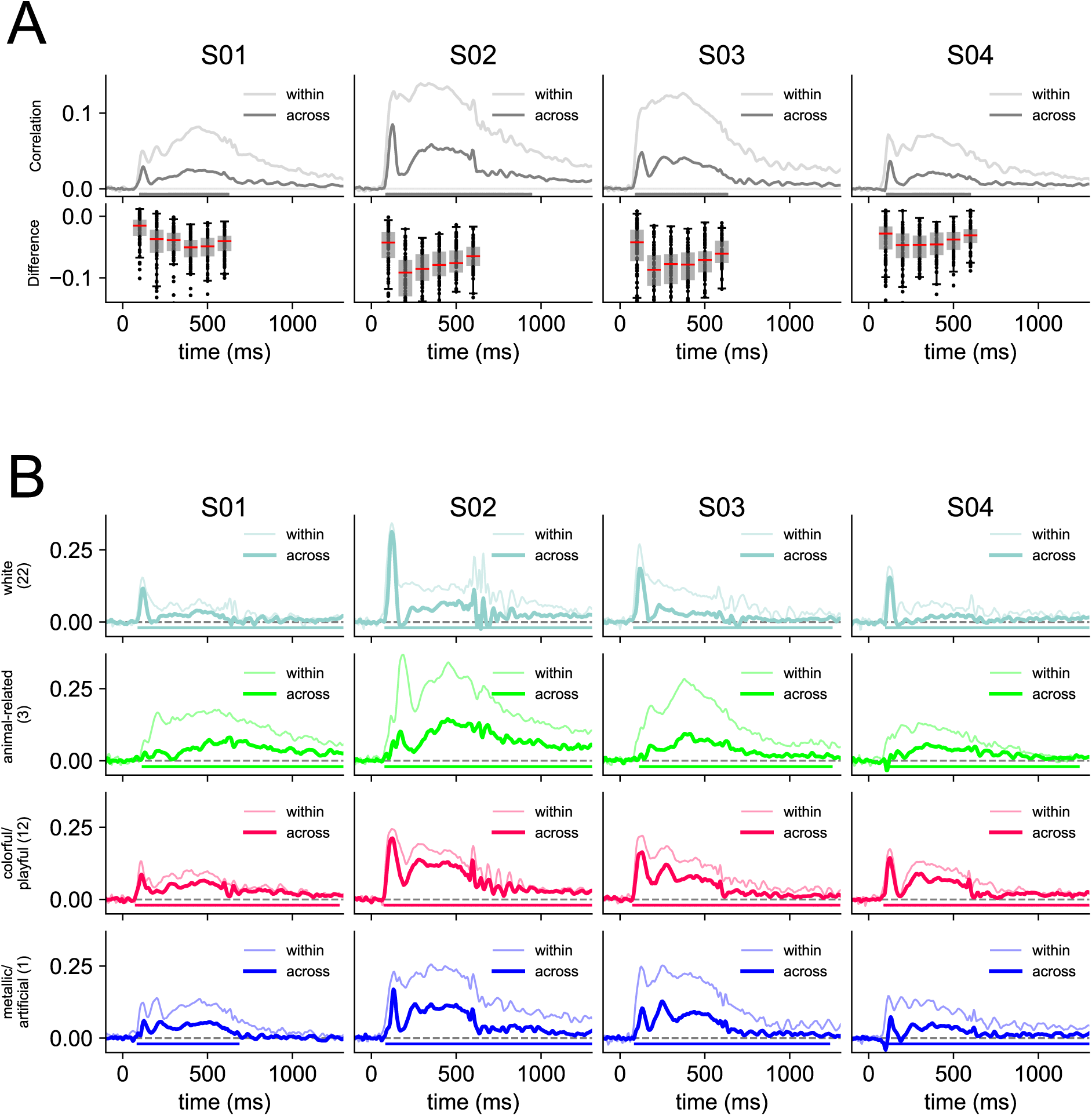
Differences between within- and across-participant model. (A) Average performance of the model across all dimensions when fitted as a within-participant model (session-wise cross-validation) and an across-participant model (participant-wise cross-validation). The dots below the time courses show significant timepoints for each of the four participants for the across-participant model at a threshold of p < 0.01 established based on null distributions generated using label permutations (see Methods). The difference between the two models is plotted at select time windows of interest at 100 ms, 200 ms, 300 ms, 400 ms, 500 ms, and 600 ms. Statistics of the pairwise comparisons for each time window are reported in supplementary material (see Figure S3). (B) Each subplot shows an example dimension timeseries when the model is fit within each participant (light color) and across different participants (dark color). All other dimension time courses can be found in the supplementary materials (see Figure S3).

### (3) Peak timing and relative amplitude are prototypical temporal characteristics of different dimension time courses

Comparing the dimension time courses visually suggests some commonalities across dimensions. For example, some dimensions shared a strong early peak and others showed a slower, gradual rise. To quantify the similarity of time course shapes across dimensions, we used dynamic-time warping (DTW, Chu et al., 2002). DTW captures the similarity between a pair of timeseries by assessing how much one of the timeseries has to be warped to resemble another one (Figure 4A). The result of this analysis is a time-time-matrix with cost values indicating the amount of warping that has to be done at every timepoint. To measure the similarity of a given timeseries pair, we extracted the sum of the Euclidean distances along the path of lowest cost. If the path falls on the diagonal of the time-time-matrix it means that the timeseries are identical. If it veers off the diagonal, the timeseries are more dissimilar (Figure 4A).

**Figure 4.**
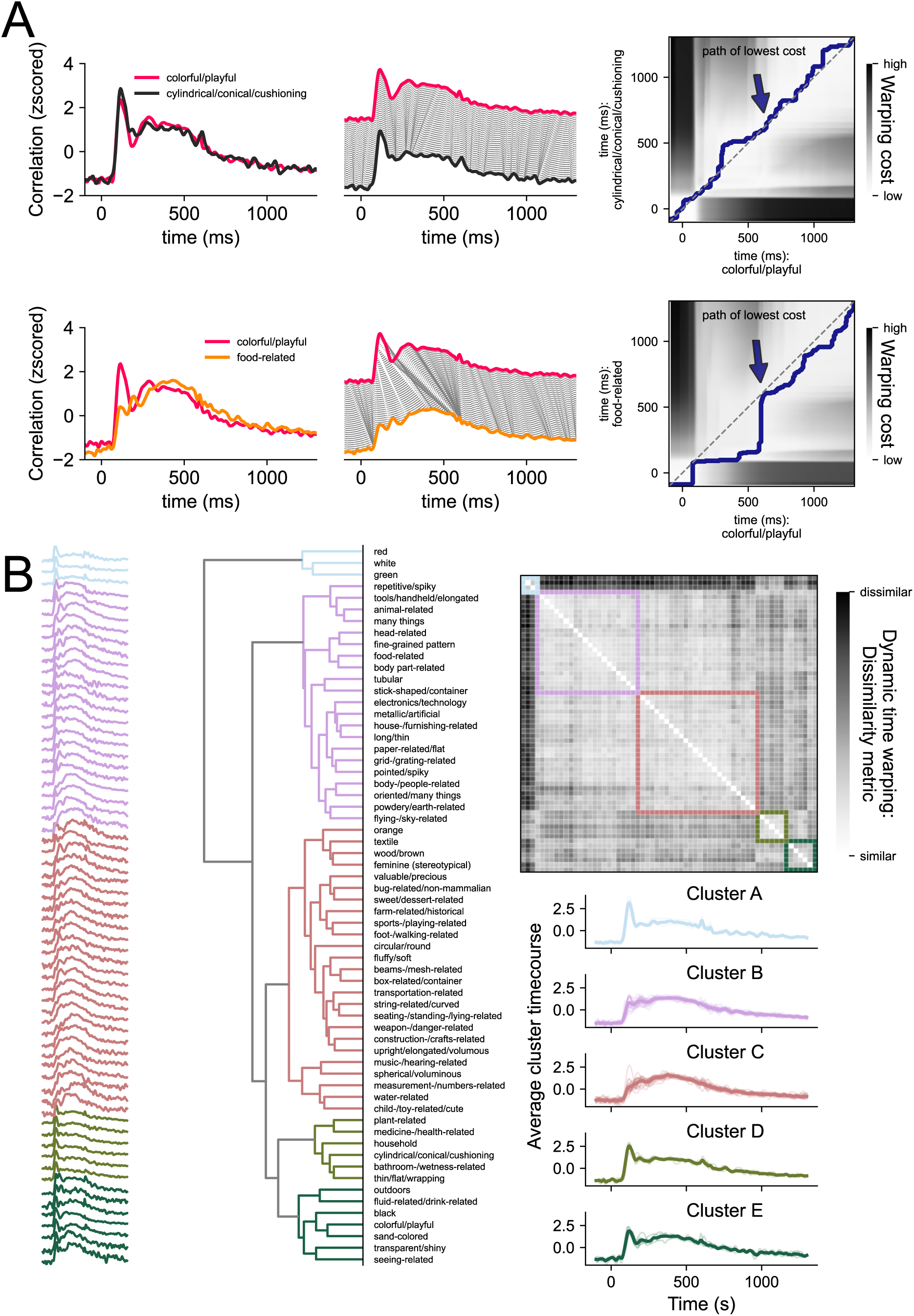
Dynamic time warping (DTW) as a method to compare timeseries similarities and extract prototypical timeseries characteristics. (A) shows the DTW approach for two pairs of timeseries. The correlations were scaled and plotted over time (left panel). DTW assesses how much one timeseries needs to be warped to resemble the other one (middle panel). This warping cost can be established by calculating the Euclidean distance between all timepoints (top right panel) to assess dissimilarity. The path of lowest cost describes the path through the matrix that minimizes the warping costs while adhering to some rules (see Methods). To summarize the warping cost in a single number, we summed the Euclidean distances along the path as a dissimilarity measure. (B) shows the dissimilarity matrix containing the DTW similarity measures for timeseries pairs (right panel). Timeseries with low signal-to-noise ratio were excluded (see methods). Hierarchical clustering on this matrix allows us to sort the dimension timeseries (left panel) and dimensions labels (middle panel). Averaging the timeseries for each cluster (bottom right panel) allows to examine prototypical timeseries characteristics.

Applying DTW to our data, we generated a distance matrix for dimension timeseries pairs and ran hierarchical clustering on that matrix to determine which dimensions evoked similar time courses (Figure 4B). Cluster A first separated from all other dimensions. This cluster contained dimensions describing colors (“red”, “white”, “green”). Next, Cluster B separated from Cluster C, which then separated from D and E. Similar to Cluster A, Clusters D and E contained dimensions describing colors (e.g., “sand-colored”, “black”), have very consistent colors (e.g., “plant-related”, “bathroom-related”) as well as other features such as shape (e.g., “cylindrical/conical/cushioning”, “thin/flat”). In contrast, Clusters B and C contained properties that are visually more varied (e.g., “animal related”, “many things”). After running the clustering, we sorted and averaged the cluster correlations to examine prototypical timeseries characteristics. The primary feature that appeared to distinguish the different time series was the relative strength of the early (∼ 125 ms) and late (> 200 ms) correlations. The presence of an early peak (clusters A, D and E) was also often accompanied by a second local peak around the time of stimulus offset. Dimensions within these three clusters included “red”, “green”, “thin/flat”, “transparent/shiny”, and “colorful/playful”. These were all dimensions that appear to reflect a specific visual feature (e.g. color, shape) contained within the images. Thus, the early peak and strong offset response might be driven by underlying visual consistencies in objects with high scores on these dimensions.

In contrast, the other clusters showed the strongest correlation after 200 ms, with a slow rise to a late and prolonged maximum, but differing in the relative size of the early and late correlations. Dimensions within these clusters included “farm-related”, “flying-related”, and “body-/people-related”. Notably, these clusters accounted for the majority of the dimensions, highlighting the importance of the later response for behaviorally relevant features.

Overall, the clustering suggests that there are two broad classes of temporal profiles, those with a distinct, early peak and a stimulus offset effect and those with a late peak, often without any clear early peak. The clusters with strong early effects, which showed better generalization across participants, tended to reflect more stimulus-specific information (e.g. green, colorful). In contrast, the clusters with strong late peaks, which showed weaker generalization, appeared to correspond to more conceptual properties, possibly reflecting a greater contribution of subject-specific information.

## Discussion

Resolving incoming visual information to make sense of our environment is a challenging task that our brain solves within just a few hundred milliseconds. Here, we used a similarity embedding based on millions of behavioral judgements to model temporally-resolved neural responses in order to understand how behavior-derived features of objects are represented in the brain over time. Using THINGS-MEG (Hebart, Contier, Teichmann et al., 2023), we found that all individual object dimensions were directly reflected in the neural response, with distinct temporal profiles for different dimensions. In particular, while some dimensions showed strong representation shortly after stimulus onset resulting in a pronounced early peak (<150 ms) in the timeseries, other dimensions showed a slower rise to a later peak (∼300 ms). For most dimensions, these early effects were highly similar across individuals allowing for across-participant generalization, while later effects were less consistent across participants. Some dimensions that are visually more homogenous (e.g., “red”) exhibited high across-participant generalization throughout time. Collectively, these results indicate that dimensions capturing physical stimulus properties evoke a more similar response across participants. Overall, our work highlights that behavior-derived object properties emerge and evolve at different timepoints in the neural signal and contribute to the rich nature of object vision.

The current study builds on prior work suggesting early visual versus later semantic processing but extends this by focusing on behavior-derived dimensions. Our approach of combining a systematically sampled stimulus space of behavior-derived with densely sampled MEG data allows us to probe time courses of object vision more deeply. We used the >27,000 images of the THINGS database which come with a behavioral embedding derived from 4.7 million similarity judgements. Using these datasets means reduced bias in stimulus selection and category assignment (Grootswagers & Robinson, 2021), as the THINGS concepts have been selected systematically (Hebart et al., 2019), and the behavioral embeddings were derived in a data-driven fashion, relying on crowdsourced similarity judgements (Hebart et al., 2020, Hebart, Contier, Teichmann et al., 2023). The images are not repeated throughout the experiment, but the breadth of sampling allows us to model the object response directly using the crowdsourced behavioral data. Previous time-resolved analyses of response to the THINGS stimuli in MEG and EEG data revealed differential neural signals evident within the first 200 ms (Gifford et al., 2022; Grootswagers et al., 2022; Hebart, Contier, Teichmann et al., 2023) that enable object decoding. As with previous work, however, such analyses do not reveal what object properties drive these effects and the degree to which the relative contribution of different properties varies over time.

Instead of analyzing image-specific effects that reflect one category or another, we modeled the neural data for all object images in terms of continuous similarity scores along 66 dimensions. That means the MEG data going into the analysis always remained the same, but the label value capturing how strongly each image is associated with the dimension at hand differed. This approach is powerful as it makes use of all the data while allowing to study many dimensions simultaneously. Furthermore, it captures the complexities of object vision where objects are associated with many properties. The data here show that information related to all 66 behavior-derived dimensions can be read-out from the MEG signal and have specific temporal profiles. Together, our results highlight that modelling neural responses using continuous behavior-derived embeddings offers a more comprehensive understanding of the visual object space than a focus on object category or individual, experimenter-selected dimensions.

We found different temporal dynamics for different object dimensions of visual object processing with a data-driven approach that broadly revealed two distinct temporal characteristics. We observe that some dimensions had a transient and distinct, early peak <150 ms and an offset response at ∼100 ms after stimulus-offset. In contrast, other dimensions lacked the early peak completely or showed it more subtly. These dimensions tended to slowly rise to a later peak (∼300 ms) that is more sustained over time. We found that early effects were more consistent across participants than later ones, suggesting that early peaks reflect stimulus-specific and later peaks subject-specific information. Indeed, when looking at which dimensions had a distinct early peak, we found that stimulus-specific visual properties such as color drove early distinctions. In contrast, later effects seemed to be more associated with concept-related properties, and critically our results suggest that the impact of such properties was variable across participants. It is important to note that the stimulus-specific effects we observed here are not tied to specific exemplars, as every unique image was shown only once, and all analyses were based on cross-exemplar generalizations. Previous work has used cross-exemplar generalization as a method to disentangle visual and conceptual object properties (e.g., Bankson, Hebart et al., 2018; Carlson et al., 2013), however, this approach does not allow us to distinguish which object properties drive the effects at different timepoints. Our approach uses multidimensional behavioral embeddings and can therefore tease these differences apart by showing which behavior-derived property contributes to the neural signal at a given timepoint. Overall, the results highlight that specific temporal profiles are associated with different behavior-derived dimensions but that some broad characteristics can distinguish between stimulus- and subject-specific information.

The results of the current study build on and advance findings from earlier studies on object vision which primarily examined broad object categories or specific features and examined how differences between those arise in the human brain (e.g., Carlson et al., 2013a; Cichy et al., 2014; Clarke et al., 2013; Goddard et al., 2016; Grootswagers et al., 2019; Hebart et al., 2018; Liu et al., 2002; Rossion, 2014; van de Nieuwenhuijzen et al., 2013). For example, prior EEG and MEG studies revealed differences in responses to faces and other objects that peak around 170 ms (Liu et al., 2002; Rossion, 2014). More recently, using multivariate and machine learning analyses, studies have further shown that M/EEG signals evoked by broad object categories (e.g., animals, plants, body parts) can be distinguished within the first 200 ms (Carlson et al., 2013; Cichy et al., 2014; Clarke et al., 2013; Goddard et al., 2016; Grootswagers et al., 2019; Hebart et al., 2018; van de Nieuwenhuijzen et al., 2013). To disentangle what features might be driving differences in the neural response, some studies have used stimulus sets with perceptually similar stimuli (e.g., glove and hand) or stimuli that straddle category bounds of object properties such as animacy (e.g., robots) (Contini et al., 2021; Kaiser et al., 2016; Proklova et al., 2019). Others have tried to separate the contribution of visual and semantic object properties to the neural signal by using cross-exemplar generalization to determine when we can distinguish objects across different exemplars (Bankson, Hebart et al., 2018; Carlson et al., 2013), across object position and size (Isik et al., 2014), or modelling the data using visual and semantic models (Clarke et al., 2015). These studies, though foundational in understanding the neural time course of object vision, often relied on small or biased stimulus sets (e.g., Bankson, Hebart et al., 2018; Carlson et al., 2013; Cichy et al., 2014; Grootswagers et al., 2018; Kriegeskorte et al., 2008; Teichmann et al., 2020) with findings that may be tied closely to the specific stimuli chosen (Grootswagers & Robinson, 2021). While larger stimulus sets potentially avoid this problem, it is often unclear how the stimuli were sampled, and biases within the stimulus set may still constrain the results. For example, the COCO stimuli used in the large-scale MRI Natural Scenes Dataset (Allen et al., 2022) heavily oversample 80 individual categories, leading to countless images of giraffes or surfers, which may also overestimate generalization performance (Shirakawa et al., 2024). Second, analyses in prior work often focused on category or feature labels assigned to individual stimuli, which ignore our broader understanding of objects and the specific properties that may be shared between different objects (Ritchie et al., 2024). One alternative approach used in prior work is to model the neural data using feature norms (e.g., McRae et al., 2005). In one particular MEG study (Clarke et al., 2015), this approach was used to model semantic content of stimuli from 11 categories which was then contrasted with output from a computational model of object vision. A drawback of this approach is that feature norms rely on verbally naming properties which means key visual or conceptual features may be missed while other features may be overemphasized. In contrast, our approach extracts object properties that are behavior-derived using a visual odd-one-out task which does not require properties to be nameable and automatically extracts their individual relevance.

One limitation of our work is that the behavioral embedding was derived from a separate set of participants than those from which neural responses were collected. While our data is consistent enough to be generalizable across participants, we find that generalization performance is better for earlier peaks and dimensions that capture perceptually homogenous features (e.g., red, green, colorful). This may partially be the case because our behavioral embeddings are derived from crowdsourced data and thus may prioritize dimensions that tend to be shared across individuals. Future work should investigate individual differences more closely to understand how the object space may be skewed given personal experience and the task at hand.

In conclusion, by using behavioral judgments of similarity to guide our understanding of the neural representation of the object space, we find that different aspects of the object response emerge at different timepoints and together create the experience of meaningful visual object processing.

## Methods

### Dataset

We used the publicly available THINGS dataset (Hebart, Contier, Teichmann, et al., 2022) which contains densely sampled MEG data as well as crowdsourced behavioral data. The MEG portion of the data contained neural recordings from four participants who each viewed a total of 27,048 unique images over the course of a 12-sessions. Every image was shown centrally for 500 ms with an inter-stimulus interval of 800-1200 ms. Participants completed a target-detection task, looking for artificially generated images of objects that do not exist. Of the 27,048 trials, 22,248 trials were experimental trials showing unique images from the THINGS database (Hebart et al., 2019), which were used for the analysis.

Each image belonged to one of 1,854 object concepts (e.g., aardvark, clock, chicken wire, among many others). Unique image exemplars for each of the concepts were repeated 12 times over the course of the MEG experiment (one image per concept per session). In addition to the image concepts, we used a behavior-derived embeddings to model MEG sensor responses. The embeddings contained weights on 66 dimensions which capture trial-by-trial responses for 4.7 million odd-one-out judgments on triplets of the 1,854 object concepts (Hebart, Contier, Teichmann et al., 2023). Each dimension describes a certain object property (e.g., circular/round, colorful, food-related), however, these dimensions were derived in a data-driven way based on the behavioral data. The original embedding was trained at the concept-level (one image per concept) and hence could miss visual variability across exemplars. In order to obtain image-level embeddings, we used a neural network model (CLIP-ViT, Radford et al., 2021) that can predict image-text pairs and has also been shown to be able to predict similarity judgments with high accuracy (Hebart et al., 2022; Muttenthaler et al., 2022). We used a separate ridge regression model for each dimension and fit it for the behavioral data corresponding to 1,854 images. Then we examined the activity patterns in the final layer of the image encoder. We then used ridge regression to predict dimension weights for each of the 66 dimensions for all ∼27,000 images in the THINGS database. To model the evoked neural response measured with MEG, we then used the image-level predicted weights along the 66 dimensions. Please note that, while this analysis relies on features derived from a neural network model, the human similarity embedding showed good fits at the level of individual dimensions, demonstrating that these effects were not merely driven by projecting the CLIP image embedding to 66 arbitrary unidimensional spaces.

### Preprocessing

Our preprocessing pipeline was built using *mne-python* (Gramfort et al., 2013) and described in detail in the dataset release (Hebart, Contier, Teichmann, et al., 2022). The preprocessing steps included filtering (1Hz – 40Hz), baseline correction using z-scoring and epoching the data from -100 to 1300 ms relative to stimulus onset. Raw and preprocessed data can be directly downloaded from OpenNeuro (https://openneuro.org/datasets/ds004212/versions/2.0.0).

### Analyses

#### Modelling MEG data based on multidimensional similarity judgments: within-participant regression

To model how the behavior-derived dimensions unfold over time in the human brain, we fitted multiple ridge regression models at every timepoint to learn the association between the multivariate MEG-sensor response and the scores along each dimension. We trained the model on data from 11 out of the 12 sessions (20,394 trials) and tested on the remaining one (1,854 trials). This process was repeated so that every session was used as testing data once. The model was trained and tested for each participant separately. A separate model was trained and tested at every timepoint. Models were fit in Python using ridge regression models with a tuned alpha regularization term at each timepoint (possible alpha regularization terms included 100 linearly spaced values between 0.000001 and 30,000). The input data was scaled using the *sci-kit learn StandardScaler* function and the regularization was tuned in an internal cross-validated way using the *sci-kit learn RidgeCV* function (Pedregosa et al., 2011).

We assessed the model’s performance by correlating the predicted dimension score of all left-out trials with behavioral embeddings for each of the images. These correlations were interpreted as amount of information in the neural signal associated with a given dimension. To assess statistical significance, we generated subject-level null distributions. We re-ran the identical analysis using a session-wise cross-validation but shuffled the testing labels 1,000 times. The shuffling index was kept consistent across cross-validation splits and the resulting correlations were averaged. Thus, for each dimension we had a null distribution at every timepoint. We defined the 99^th^ percentile of each distribution as a threshold and then looked for the maximum value across timepoints and dimensions to test the model performance against to correct for multiple comparisons.

To gain insights into which sensors primarily drove the effects, we also trained a linear model to predict the activation of each sensor using the multidimensional similarity judgments. We used a session-wise cross-validation approach and ran this analysis for each participant separately. The model’s performance was assessed by correlating the predicted sensor activations for the test-set and the true sensor activations at every timepoint.

### (b) Examining time course similarities across people: Across-participant regression

To examine whether time course profiles are consistent across participants, we trained the a model to learn the association between the multivariate MEG sensor activation pattern at every timepoint and the behavioral dimension profiles using data from one of the participants and testing it on the others. We did a session-wise cross-validation to match the within-participant analysis in terms of training and testing set size (see (a)). We repeated this process until all pairwise comparisons of participants were used as training and testing set. A separate model was trained and tested at every timepoint. Models were fit in Python using *sci-kit learn* ridge regression models following the same hyperparameter tuning method as in the within analysis.

We plotted the difference betwene the within- and the across-participant model for all 66 dimnesions at select timepoints (100 ms, 200 ms, 300 ms, 400 ms, 500 ms, 600 ms). We used bonferroni-corrected pairwise t-tests to test whether the differences between the models were significant (defined as p < 0.01) at these timepoints. The results of the tests are reported in the supplementary materials (Figure S3).

### (c) Examining time course similarities across dimensions: Dynamic Time Warping

The results of the regression models were timeseries of correlations for each dimension. To compare the shapes of these timeseries and assess overall similarities and differences, we used dynamic time warping implemented in the *dtaisdistance* toolbox (Meert et al., 2020). In contrast to correlation measures which are compression based, DTW is shape based and is well suited to investigate timeseries similarities that may have a temporal drift (Aghabozorgi et al., 2015). The goal of DTW is to find matches between patterns of two timeseries by assessing how much one timeseries has to be warped to look like the other one. This is achieved by generating distance matrices filled with pairwise Euclidean distances between timepoints and finding the shortest path through this matrix while adhering to several rules: The start and end of the timeseries have to align, the path cannot go back in time, and it has to be continuous.

The DTW similarity measure represents the sum of the Euclidean distances along the shortest path. We extracted this measure for smoothed time courses and generated a similarity matrix. Given that we were interested in the relative shape of the timeseries and not the differences in signal-to-noise ratio, we normalized the timeseries before running the dynamic time warping by calculating z-scores for each dimension timeseries and each participant. Then we averaged across participants and calculated the DTW similarity measure for all dimension comparisons. Because z-scoring and timeseries with only a few timepoints that are above zero can amplify the effect of noisy timeseries, we sorted the time courses by number of significant timepoints across all participants and excluded the dimension time courses with the lowest 5% of significant timepoints. This resulted in three dimension time courses to be excluded (fire-related, yellow, masculine (stereotypical)). As this cutoff is arbitrary, we also tested to exclude more dimensions or basing the exclusion on a different metric (e.g., peak amplitude before z-scoring). The bottom dimensions are always the same and the number of excluded dimensions does not influence the conclusions from the DTW analysis in any substantial way.

Running hierarchical clustering on the resulting DTW distance matrix allowed us to establish a qualitative measure of different prototypical timeseries characteristics. We set the threshold for the hierarchical clusters to be at 0.5 x the maximum distance observed. This threshold is arbitrary, and the number of clusters extracted can be influenced by changing this threshold. However, the overall conclusion that the two main characteristics differentiating the clusters are an early and sharp peak versus a slow rise to a late, sustained peak are robust.

### Open Science Practices

All data is publicly available under a Creative Commons license and can be downloaded from OpenNeuro: https://openneuro.org/datasets/ds004212/versions/2.0.0 *[this repository does not contain all aggregate behavioral data to run the analyses yet. It will be populated and re-uploaded as version 3.0.0 as soon as the review process is finalized].* The analysis code for all analyses in this paper are available on GitHub: https://github.com/Section-on-Learning-and-Plasticity/THINGS-MEG.

## Supplementary Materials

**Figure S1.**
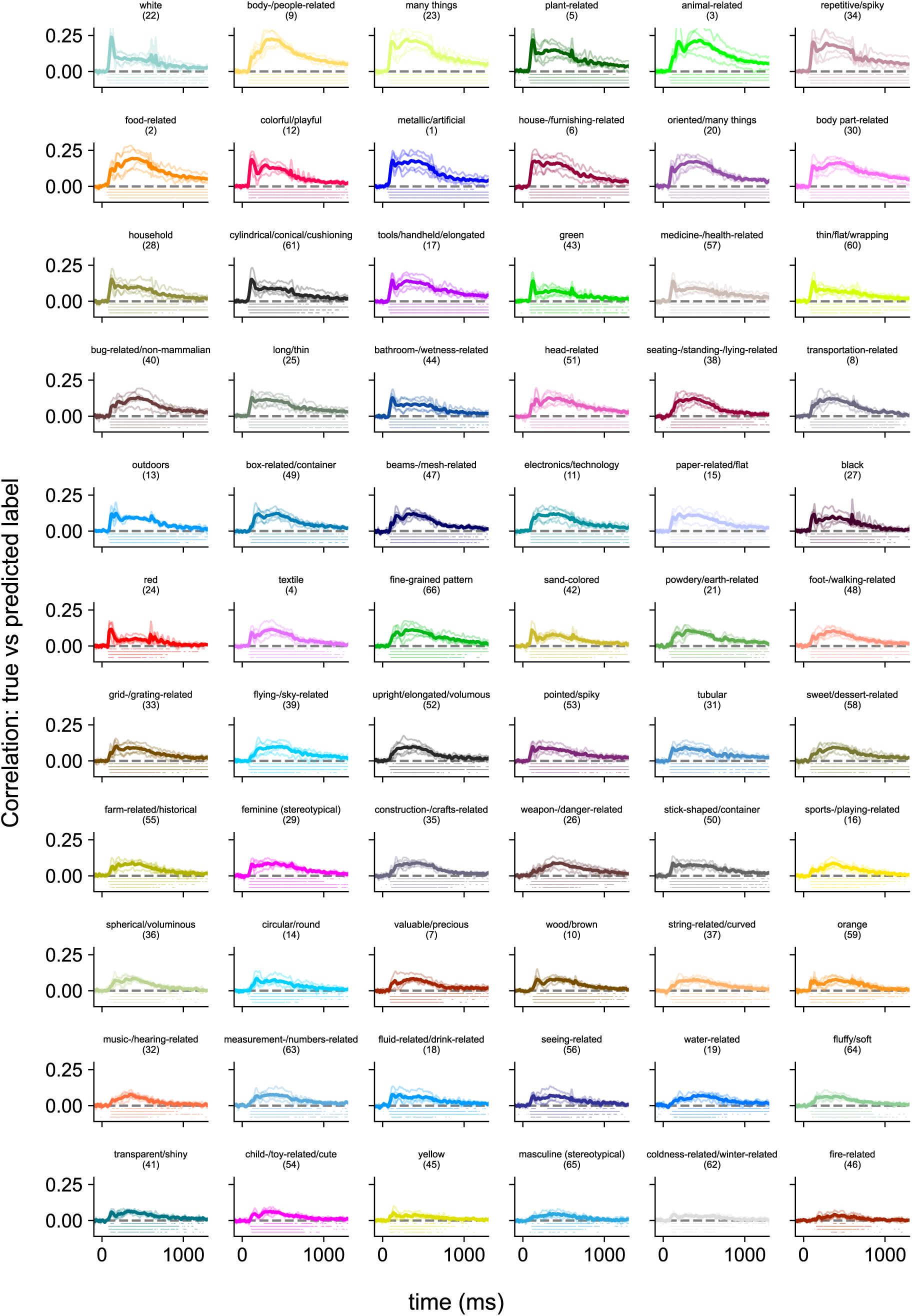
Dimension time courses for within-participant regression model. Each panel shows the correlation between the predicted and true weights for each dimension over time. The time courses are sorted by peak amplitude. The thin lines show the correlations for each participant. The thick lines show the average across participants. Dots below the timeseries show significant timepoints at p < 0.01 based on subject-level null distributions derived from permutations (see Methods).

**Figure S2.**
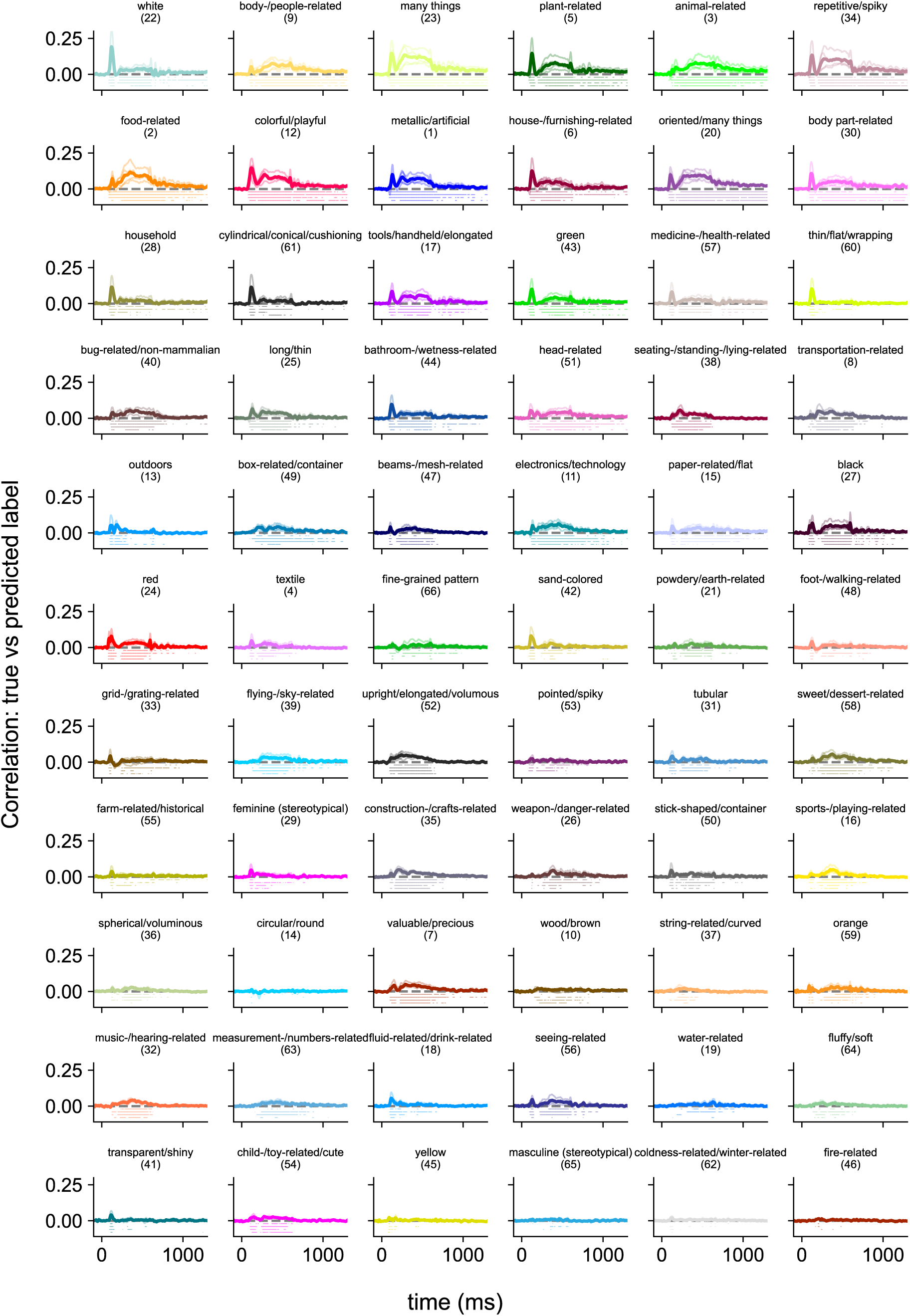
Dimension time courses for across-participant regression model. Each panel shows the correlation between the predicted and true weights for one of the 66 dimensions over time when the model is trained and tested on data from different participants. The thin lines show the correlations for each participant. The thick lines show the average across participants. Dots below the timeseries show significant timepoints at p < 0.01 based on subject-level null distributions derived from permutations (see Methods).

**Figure S3.**
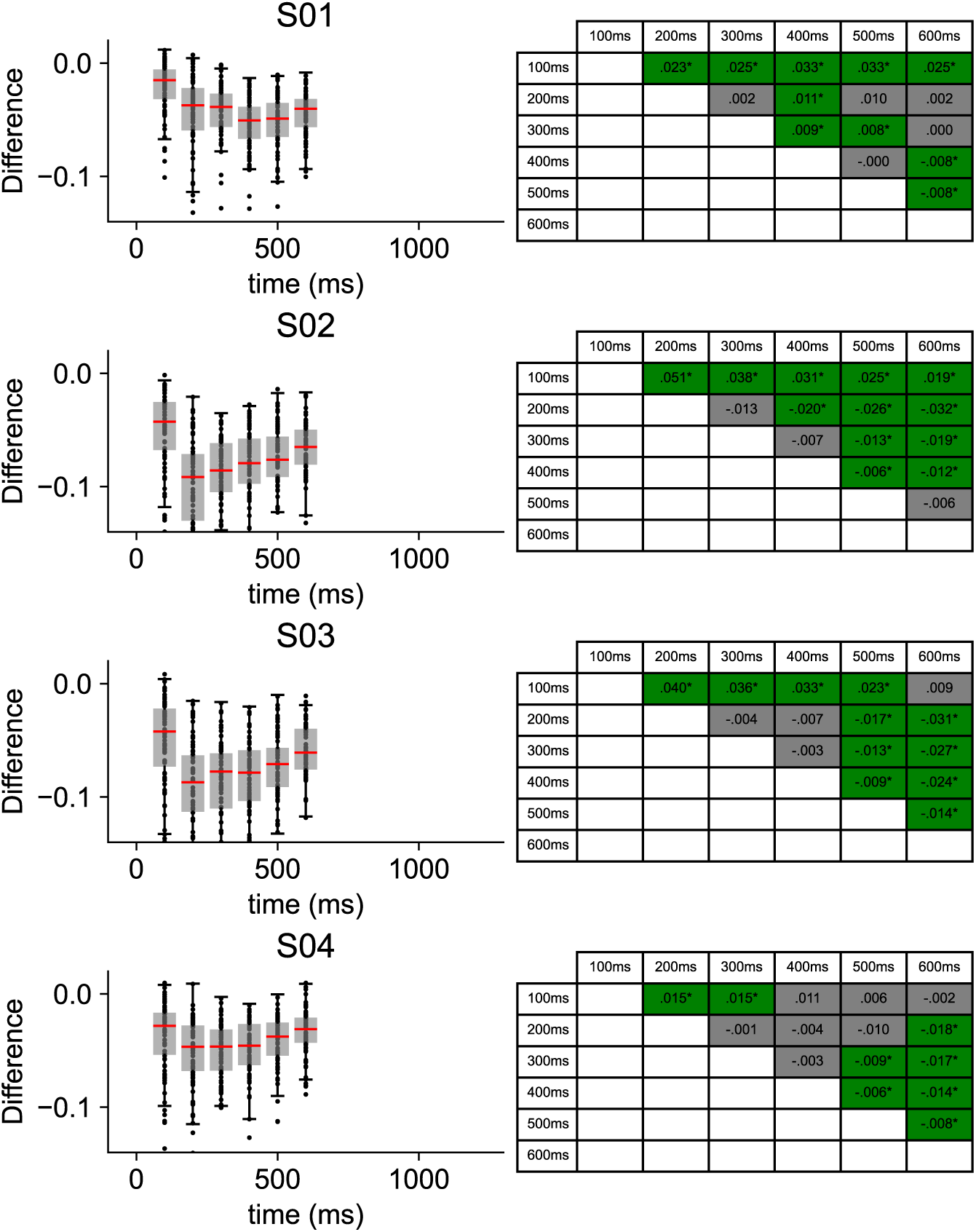
Differences in within- and across-participant models. Each row shows the data from one participant. The left panels show the difference between the within and the across participant model across the 66 dimensions at 6 different timepoints of interest. The right table shows the differences between each pairwise comparison of the timepoints of interest with green boxes (and stars) highlighting whether the comparison is significantly different from zero (Bonferroni corrected, p < 0.01).

## Acknowledgements

We thank Anna Corriveau, Alexis Kidder, Adam Rockter, and Maryam Vaziri-Pashkam for their help collecting the data and Philipp Kaniuth for help generating the predicted similarity embedding. Additional thanks to Tom Holroyd and Jeff Stout for technical support and discussions. We thank Grace Edwards and Susan Wardle for helpful comments on earlier versions of this manuscript. We utilized the computational resources of the NIH HPC Biowulf cluster to run the MEG analyses (http://hpc.nih.gov). The work presented here was supported by the Intramural Research Program of the National Institutes of Health (ZIA-MH-002909), under National Institute of Mental Health Clinical Study Protocol 93 M-1070 (NCT00001360). Additional support was received by a research group grant by the Max Planck Society awarded to M.N.H., the ERC Starting Grant project COREDIM (ERC-StG-2021-101039712), and the Hessian Ministry of Higher Education, Science, Research, and the Arts (LOEWE Start Professorship to M.N.H. and Excellence Program “The Adaptive Mind”)

